# Monsters in the uterus: A parthenogenetic quasi-program causes teratoma-like tumors during aging in wild-type *C.elegans*

**DOI:** 10.1101/174771

**Authors:** Hongyuan Wang, Yuan Zhao, Marina Ezcurra, Ann F. Gilliat, Josephine Hellberg, Alexandre Benedetto, Trin Athigapanich, Johannes Girstmair, Max Telford, Zhizhou Zhang, David Gems

## Abstract

Many diseases whose frequency increases with advancing age are caused by aging (senescence), but the mechanisms of senescence remain poorly understood. According to G.C. Williams and M.V. Blagosklonny, a major etiological determinant of senescence is late-life, wild-type gene action and non-adaptive execution of biological programs (or quasi-programs). These generate a wide range of senescent pathologies causing illness and death. Here we investigate the etiology of a prominent senescent pathology in the nematode *C. elegans*, uterine tumors, in the light of the Williams Blagosklonny theory. Uterine tumors develop from unfertilized, immature oocytes which execute incomplete embryogenetic programs. This includes extensive endomitosis, leading to formation of chromatin masses and cellular hypertrophy. The starting point of pathogenesis is exhaustion of sperm stocks. The timing of this transition between program and quasi-program can be altered by blocking sperm production (causing earlier tumors) or supplying additional sperm by mating (delaying tumor onset). Other pathophysiological determinants are yolk consumption by tumors, and bacterial proliferation within tumors. Uterine tumors resemble mammalian ovarian teratomas (*tera*, Greek: monster) in that both develop from oocytes that fail to mature after meiosis I, and both are the result of quasi-programs. Moreover, older but not younger uterine tumors show expression of markers of later embryogenesis, i.e. are teratoma-like. These results show how uterine tumors in *C. elegans* form as the result of run-on of embryogenetic quasi-programs. They also suggest fundamental etiological equivalence between teratoma and some forms of senescent pathology, insofar as both are caused by quasi-programs.

## Introduction

Thanks to advances in treatments for infectious diseases, aging (i.e. senescence) has now become the main cause of mortal disease worldwide. However, the causes (etiologies) of aging remain poorly understood. One approach to investigate aging is to employ simple, experimentally tractable model organisms such as the nematode *Caenorhabditis elegans*. Since the 1990s, major advances have been made in terms of identifying genes and pathways with effects on *C. elegans* lifespan, but less so in terms of understanding the underlying processes of aging that such genes influence.

The presence of so many genes where loss of function increases lifespan implies that wild-type gene action is a major cause of aging. This is consistent with G.C. Williams' evolutionary principle of antagonistic pleiotropy (AP): that natural selection can favour alleles that enhance fitness in early life even if they promote pathology later in life. This can occur because natural selection declines with age after the onset of reproduction (Abrams, 1993; Williams, 1957).

Critical to understanding aging is identification of the mechanisms by which AP is enacted, and there are several theories about this. The disposable soma theory suggests that late-life costs reflect insufficient resource investment into somatic maintenance mechanisms that protect against molecular damage that causes aging (Kirkwood, 1977). An alternative model, proposed by M.V. Blagosklonny, argues that the predominant cause of life-limiting senescent pathology is not molecular damage, but rather late-life wild-type gene action (or *hyperfunction*), which leads to pathogenic activation of biological programs (developmental, reproductive, reparative) (Blagosklonny, 2006; Blagosklonny, 2008). Such functions are programmed by genes yet not programmed in the sense of being an adaptation, i.e. are quasi-programmed. By this view, senescence originates with the transition from program to quasi-program. Mammalian examples of quasi-programmed processes include the promotion of cancer in late life by senescent cells (sensu Hayflick) (Campisi, 2013), or the promotion of atherotic plaque formation by inflammation.

Here we use the Williams Blagosklonny framework of ideas to investigate aging in *C. elegans*. Accordingly, we focus on understanding how wild-type gene action generates pathology. This involves a change in approach from what is traditional in *C. elegans* biogerontology, in that we use senescent pathology rather than lifespan as a metric of aging: ultimately, discovering the origins of senescent pathologies is key to understanding aging. Our approach involves a plausible assumption: that when *C. elegans* die of old age, it is as the result of senescent pathologies. Moreover, not all senescent pathology is life-limiting, and the identity of life-limiting pathologies will vary between individuals, culture conditions and species. By this view, understanding the determinants of lifespan in *C. elegans* are less important, in terms of applicability to human aging, than understanding the causes of senescent pathology.

Compared to the genetics of lifespan, the biology of *C. elegans* senescent pathology has been relatively neglected, apart from a few pioneering studies e.g. from the Kenyon, Driscoll and Melov labs (Garigan et al., 2002; Golden et al., 2007; Herndon et al., 2002; McGee et al., 2012; McGee et al., 2011). Wild-type hermaphrodites exhibit a range of senescent pathologies whose severity and rapidity of development suggests the action of quasi-programs (Gems and de la Guardia, 2013), and which involve atrophy (e.g. intestine, body wall muscles and gonad) and hypertrophy (e.g. uterus, some neurons and body cavity steatosis). The most striking hypertrophic pathology in aging hermaphrodites, seen in all individuals, are the uterine tumors (also referred to as tumor-like masses or oocyte clusters) (Hughes et al., 2011; Luo et al., 2010; Riesen et al., 2014). These arise from the germline of *C. elegans* hermaphrodites, which initially generates sperm that are stored in the spermatheca, but then switches to oocyte production (protandry). Oocytes are fertilized as they pass through the spermatheca, until sperm stocks are depleted. After that, oocyte production continues for a while, and oocytes pass unfertilized from the spermatheca into the uterus. Many such oocytes are expelled through the vulva, but a few remain in the uterus where they undergo a striking transformation, undergoing major cellular hypertrophy to form tumors so large that they can fill the entire body cavity in the *C. elegans* mid-body (Figure 1A). Despite their massive size (relatively speaking), uterine tumors do not contribute to age-related mortality in wild-type *C. elegans* under standard culture conditions: preventing their development by blocking germline development or through use of an inhibitor of DNA replication (5-fluoro-deoxyuridine, FUDR) does not increase lifespan (Hsin and Kenyon, 1999; Riesen et al., 2014). FUDR (Floxuridine) is a widely used treatment for colorectal cancer, a major pathology of the elderly and the third leading cause of cancer death in the U.S. (Jemal et al., 2002): this illustrates the value of investigating the etiologies of senescent pathology in *C. elegans* regardless of their effects on nematode lifespan.

**Figure 1.**
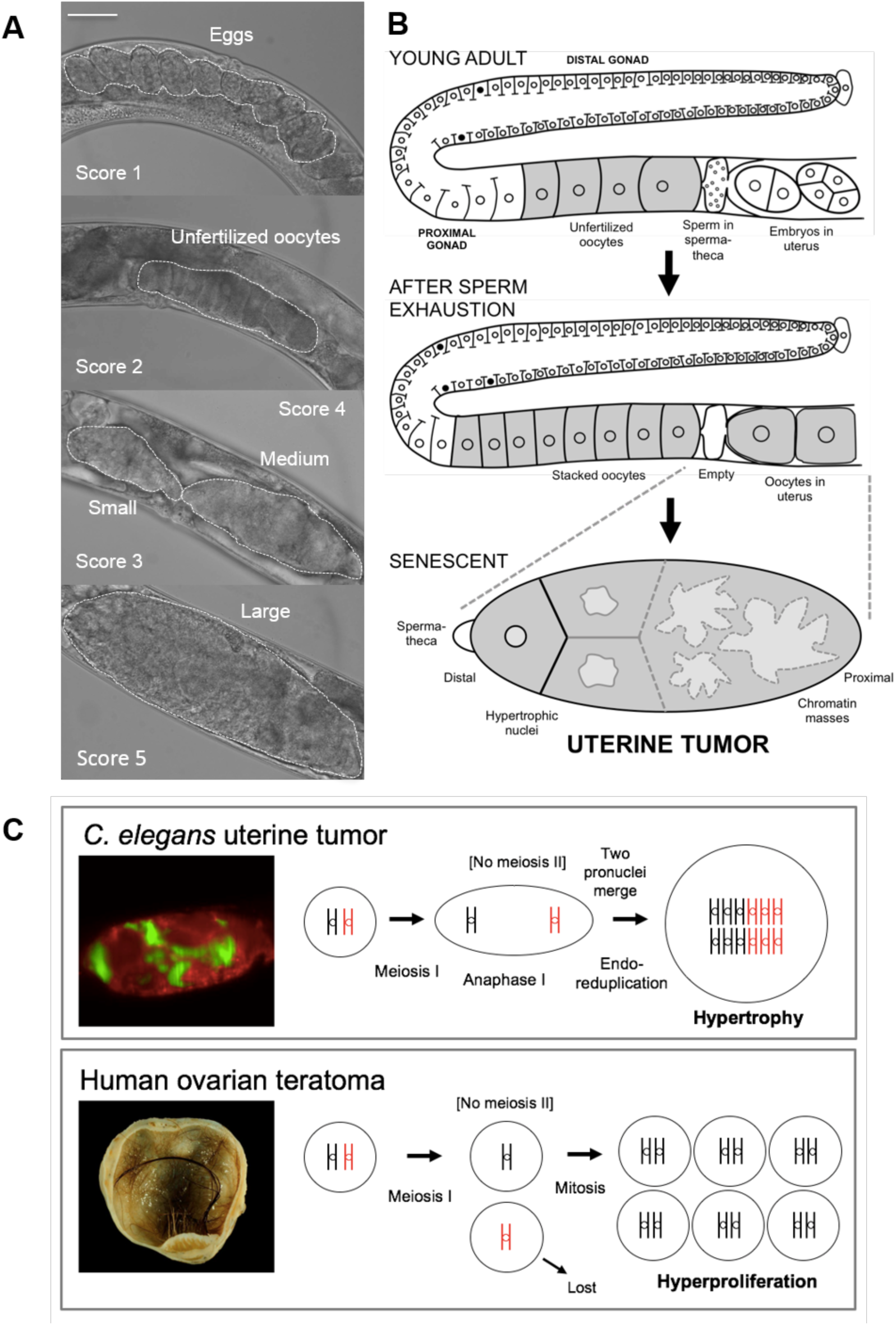
Characteristics of senescent uterine tumors. (A) Stages of development of uterine tumors (Nomarski microscopy). Scale bar, 50 μm. (B) Schematic representation of uterine tumor development. The point of origin of pathophysiology is immediately after fertilization with the last sperm in the spermatheca. (C) *C. elegans* uterine tumors and human ovarian teratomas are etiologically similar in that both originate from action in unfertilized immature oocytes of quasi-programs initiated after failure of meiosis II.

*C. elegans* uterine tumors appear to result from quasi-programs. It has long been observed that endomitosis is the "normal outcome" of entry of unfertilized oocytes into the uterus after sperm depletion (Iwasaki et al., 1996; Ward and Carrel, 1979). In such oocytes there occur multiple rounds of DNA replication and nuclear envelope breakdown (M-phase entry) but no karyokinesis or cytokinesis. This is because the mitotic centriole, normally supplied by sperm, is missing (Albertson, 1984; Iwasaki et al., 1996). Such cell cycle run-on leads to nuclear hypertrophy and polyploidy (Chatterjee et al., 2005) and, it has been postulated, eventually to uterine tumors (Golden et al., 2007; Hughes et al., 2011; McGee et al., 2012) (Figure 1B).

This implies that the causes of *C. elegans* uterine tumors are similar to those of mammalian ovarian teratomas. In humans, 10-20% of abnormal ovarian neoplasms are teratomas, benign tumors forming as the result of initiation of embryonic programs in unfertilized oocytes. Within such tumors growth and differentiation can give rise to multiple tissue types and complex structures, most commonly skin, hair, muscle and cartilage (Ulbright, 2005) (Figure 1C). Other forms of teratoma can form more complex structures including bone, teeth and even partial fetuses; the grotesque appearance of some teratomas accounts for their name (*tera*, Greek: monster). Ovarian teratomas arise from a parthenogenetic quasi-program, in which maturing oocytes complete meiosis I but do not undergo meiosis II, and are thus diploid but homozygous at most loci (Eppig et al., 1977; Linder et al., 1975). In *C. elegans* the final stages of oocyte maturation, including completion of meiosis I and II, occur after fertilization, but in oocytes that remain unfertilized meiosis ceases after anaphase I (McNally and McNally, 2005). Thus *C. elegans* uterine tumors develop from diploid immature oocytes (Figure 1C).

The Williams Blagosklonny theory implies that senescent pathogenesis can, in many cases, be understood as a malign developmental process - or to use Blagosklonny's description: "twisted growth" (Blagosklonny, 2008). Thus, careful documentation of how pathologies develop is critical to understanding their origins. Here we apply this *developmental pathology* approach to better understand the pathophysiology of uterine tumors. In particular, we ask: are they the result of a parthenogenetic quasi-program? And: are they teratomas?

## Results

### Characteristics of uterine tumors

The occurrence of uterine tumors in aging wild-type *C. elegans* is well-documented (Golden et al., 2007; Hughes et al., 2011; Luo et al., 2010; Riesen et al., 2014). To better characterize this form of senescent pathology we examined tumor development using Nomarski and epifluorescence microscopy. We first asked: given that uterine tumors grow to such a great size: are they really contained wholly within the uterus? The decrepitude of older hermaphrodites can impede identification of anatomical structure, so we used a *syIs63[cog-1::GFP + unc-119(+)]* reporter strain (PS3662) in which GFP is expressed in the spermathecal-uterine valve, which marks the distal end of the uterus (and also the vulva) (Palmer et al., 2002). Tumors of 28 worms were examined on day 8 of adulthood and in all cases where spermathecal GFP was detectable, this marked the distal end of the tumor (Figure 1—figure supplement 1A,B). Thus, the tumors are contained within the uterus, which must expand as the neoplasm grows. That extra-uterine oocytes do not develop into tumors suggests the possibility that the uterus possesses tumor niche properties that are latent prior to sperm depletion (McGovern et al., 2009).

Uterine tumors are usually paired, with one developing at each end of the uterus, though larger tumors can fill the entire uterus. Anterior tumors are not larger or smaller overall than posterior ones (Figure 1—figure supplement 1C), and tumor pairs within the same worm are typically similar in size (Figure 1—figure supplement 1D). However, marked tumor size asymmetry occurs in a small fraction of worms (< 5%) and, interestingly, the majority (∼80%) of those animals also show corresponding asymmetry in the degree of gonad atrophy (Figure 1—figure supplement 1E,F). This suggests the presence of a common etiology promoting uterine tumors and distal gonad atrophy; one possibility is that resources released by distal gonad atrophy support tumor growth, another that signals from the tumor promote atrophy of the attached distal gonad.

Some tumors contained fertilized eggs embedded within them. In some cases, such embedded embryos appeared to be developing normally, but others were degenerated (Figure 1—figure supplement 1G). Embedded embryos were rarely seen in tumors of older animals. We also noted the presence of red autofluorescence within the tumors of older worms that was not present in earlier stage tumors (Figure 1—figure supplement 2A,B). Such red autofluorescence occurred in small patches at variable positions within the tumor. Notably, red autofluorescence is a biomarker of senescence in *C. elegans* (Coburn et al., 2013; Pincus et al., 2016). A possibility is that this biomarker tracks tumor patho-development, at least in part.

### Development of nuclear hypertrophy in uterine tumors

It is difficult to study tumor development using light microscopy, particularly in its later stages, due to their optical opacity and disorganized structure; they typically appear as paired, dark ovate lumps within the animal mid-body (Luo et al., 2010; Riesen et al., 2014) or as blue fluorescent masses in animals stained with the DNA-binding dye 4′,6′-diamidino-2-phenylindole hydrochloride (DAPI) (Golden et al., 2007). To view the internal structure of uterine tumors we combined two transgene arrays expressing fluorescent reporters. First, *ruIs32* which uses the *pie-1* promoter for germline expression of a GFP-tagged histone (HIS-58)(Praitis et al., 2001); this localizes to chromosomes, providing a green fluorescent marker for nuclei and DNA masses. DAPI staining confirmed that HIS-58::GFP fluorescence closely corresponds to nuclear DNA content and the presence of DNA masses (Figure 2A,B) (*p* <0.0001, linear regression analysis). Second, *ltIs44* which expresses mCherry fused to a plecstrin homology (PH) domain, also from the *pie-1* promoter (Kachur et al., 2008). PH domains bind to phosphatidylinositol 4,5-bisphosphate in the cell membrane, so mCherry:PH provides a red fluorescent marker to distinguish individual cells within tumors.

**Figure 2.**
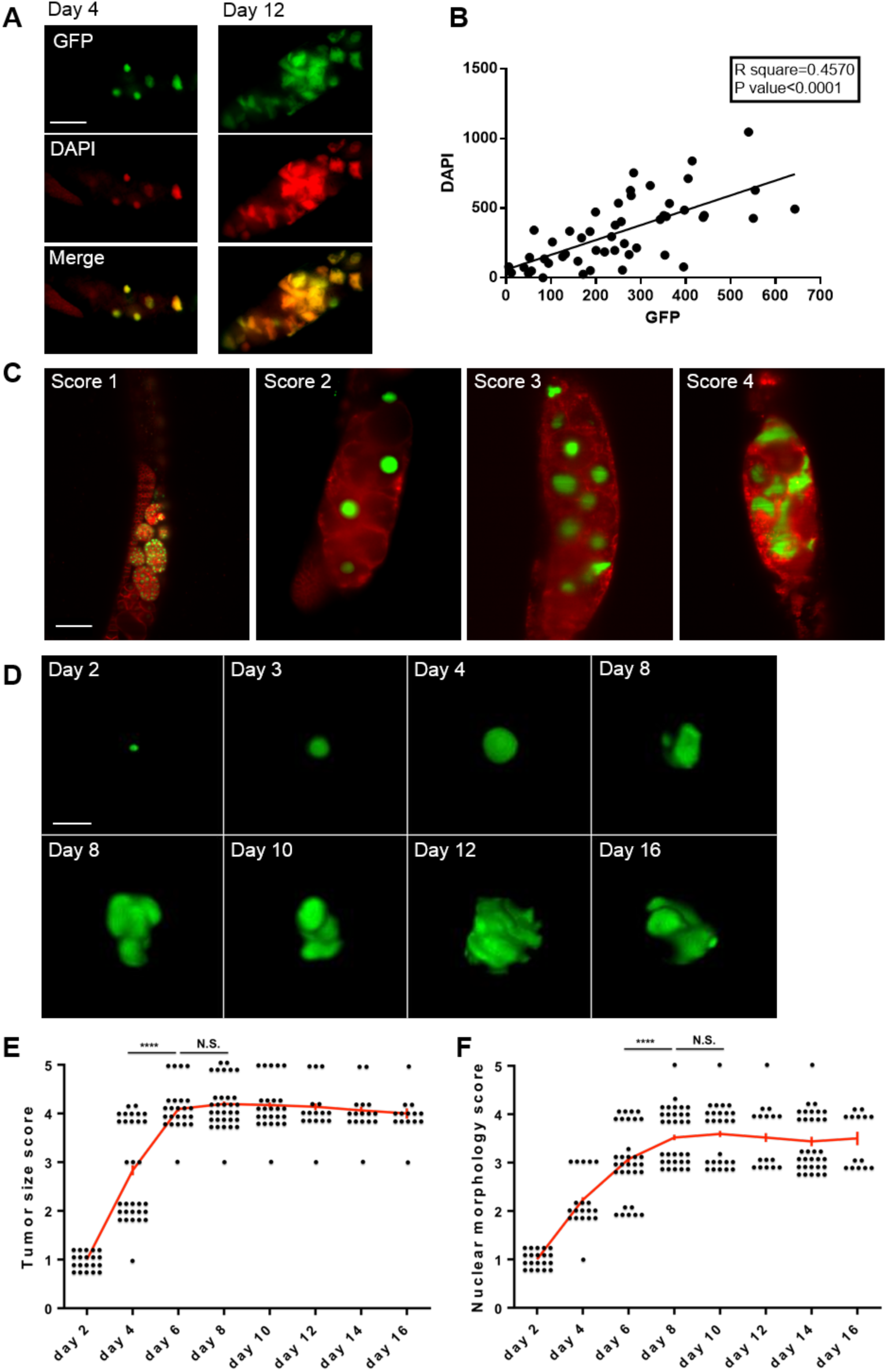
Characteristics of senescent uterine tumors. (A,B) Correspondence of HIS-58::GFP fluorescence and DNA content. (A) Tumor showing HIS-58::GFP fluorescence (top), DAPI fluorescence (center), merged image (bottom). Scale bar, 50 μm. (B) Correlation between HIS-58::GFP and DAPI fluorescence. Linear regression test, *p*<0.0001. (C) Different stages of nuclear hypertrophy of tumor. Scale bar, 25 μm. (D) Development of nuclear hypertrophy. Scale bar, 15 μm. (E) Tumor size growth during aging. (F) Changes in nuclear morphology during tumor aging. E, F, Wilcoxon-Mann Whitney test, ****, *p*<0.0001, N.S., not statistically significant.

Using the dual marker strain we examined the pattern of tumor development. In particular we wished to characterize the development of nuclear hypertrophy, and its relationship to cellular hypertrophy, and the nature of the transition from orderly spherical nuclei to disordered DNA masses. For this we used either a standard epifluorescence microscope, or confocal microscopy, or selective plane illumination microscopy (SPIM). Casual observation of SPIM images confirmed the occurrence of marked nuclear hypertrophy with increasing age (Figure 2C,D).

We then used a semi-quantitative approach to characterize the dynamics of such hypertrophy, scoring nuclear status from 1 (normal sized, spherical nucleus) to 5 (nuclei merged into large masses of chromatin)(Figure 2C). Individual worms were followed and serially photographed on alternate days, noting tumor size, and nuclear size and morphology. The main period of tumor growth was between days 2 and 6 (Figure 2E), as previously described (Riesen et al., 2014). During the initial phase of nuclear growth, between day 2 to 4, the nuclear morphology remained normal (spherical), but from around day 6 it became progressively irregular in appearance (Figure 2D,F). With increasing hypertrophy nuclear masses developed protrusions and then large, branching extensions (Figure 2D,F), which by day 8 became so large, extensive and random in structure that it became difficult to distinguish individual nuclei, which merged into amorphous masses of chromatin (Figure 2A,C,D; see videos 1 and 2 for 3D rendering of changes in nuclear morphology). There is a significant correlation between nuclear morphology and tumor size (Figure 2—table supplement 1), which may reflect the correspondence between nuclear size and morphological complexity. Although tumors reach a maximal size by day 6, nuclear masses continue to increase in size and morphological complexity until day 8 (Figure 2E,F). We also compared the number of oocytes in the tumor with tumor size on day 6 and, again, detected a positive correlation (Figure 2—figure supplement 1A).

Nuclear hypertrophy was initially more marked at the proximal end of the tumor, nearest the vulva, consistent with the greater age and more advanced patho-developmental stage of proximal cells. This proximal-distal gradient in hypertrophy disappeared in mature tumors (Figure 2—figure supplement 1B,C) as all nuclei approached their upper limit of hypertrophy.

The large increase in DNA content in aging *C. elegans* was previously linked to the appearance of DNA masses in the worm mid-body (Golden et al., 2007; McGee et al., 2012), and we verified the occurrence of this increase using PCR (Figure 2—figure supplement 1D). Comparison of the timing of age increases in intra-uterine chromatin and overall DNA content showed an overall correspondence (Figure 2—figure supplement 1E). The initial decline in DNA content could reflect disappearance of embryos, and the late decline in GFP fluorescence might be attributable to fluorophore breakdown.

### Expression of embryonic genes in mature uterine tumors

In mammalian ovarian teratomas expression of embryogenetic quasi-programs leads to pathological embryonic differentiation and morphogenesis. We wondered whether anything similar occurs within *C. elegans* uterine tumors. To this end, we first examined expression of five genes expressed in various embryonic and adult tissues but not in oocytes, *elt-1* (sperm-producing germ line)*, elt-2* (intestinal expression)*, hlh-1* (muscle)*, pha-4* (pharyngeal and intestinal expression) and *unc-119* (pan-neuronal), using GFP or EGFP reporters (Ciosk et al., 2006; Sarov et al., 2006; Zhong et al., 2010), in tumors on days 2, 4, 6, 8 and 13 of adulthood. A previous study saw no muscle or neuronal reporter expression in wild-type, polyploid unfertilized oocytes from d2-2.5 day old adults (Ciosk et al., 2006). Consistent with this, we saw no reporter gene expression in the early stages of tumor development. However, in mature tumors, after chromatin mass formation, expression of all reporters was seen, *unc-119* and *pha-4* from day 4 and *elt-1, elt-2* and *hlh-1* from day 8 or 13 (Figure 3A,B). Expression was variable, occurring in some tumors but not others, and varying in intensity in different regions of expression-positive tumors.

**Figure 3.**
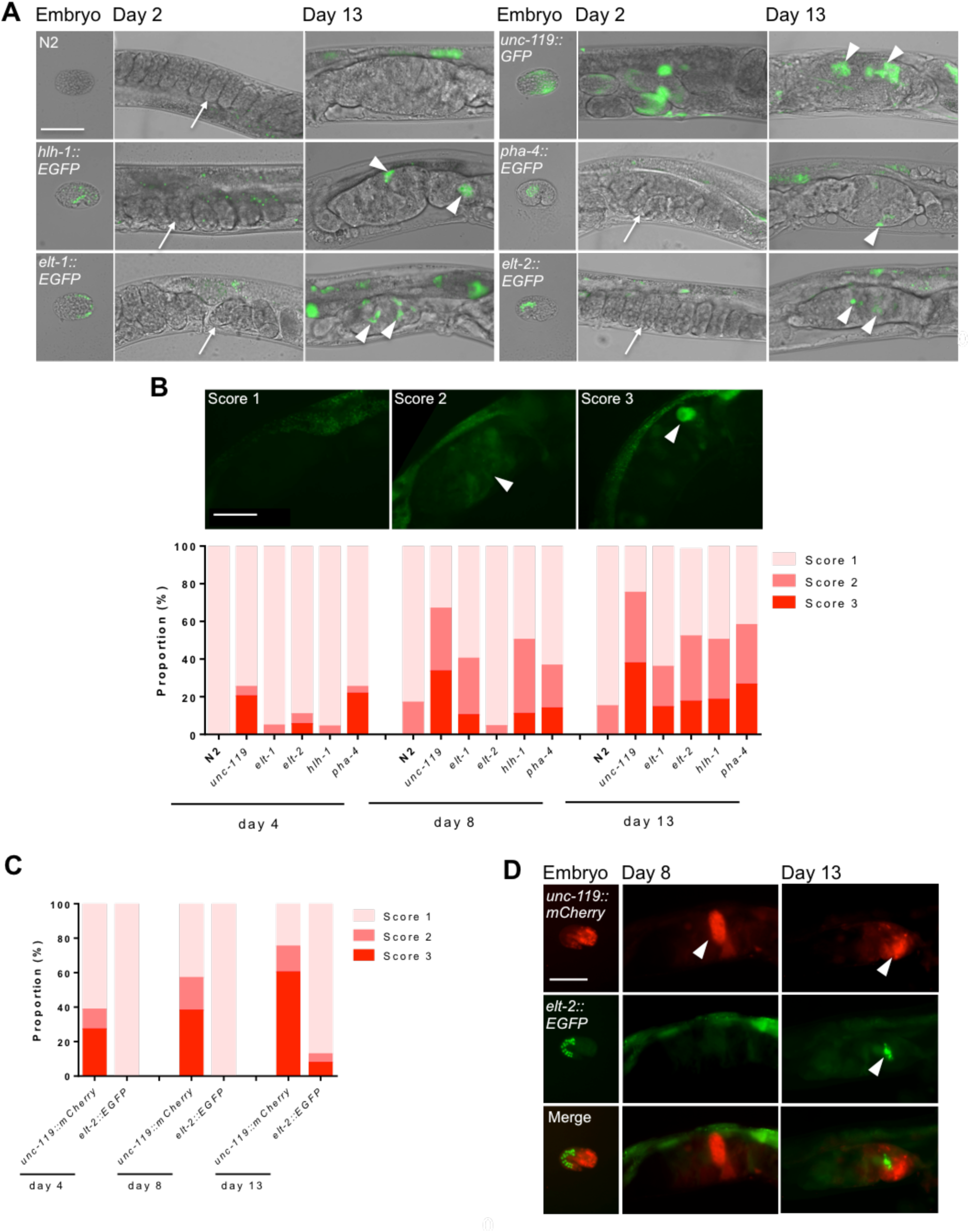
(A, B) Expression of embryonic reporters in mature uterine tumors. (A) Selected images of early and late stage tumors (epifluorescence microscopy). Note no fluorescence in early embryos (arrows). Expression was detected in older tumors (arrowheads). (B) Frequency of tumors expressing fluorescent markers and different levels (1-3 scale). Score of reporter fluorescence within tumor: score 1, no fluorescence; score 2, weak fluorescence; score 3, strong fluorescence. GFP fluorescence in uterine tumors (arrowheads: fluorescence). (C, D) Expression within the same tumor of *unc-119::GFP* and *elt-2::EGFP*. (C) Frequency of tumors expressing fluorescent markers and different levels. (D) Example of day 13 tumor expressing both *unc-119::GFP* and *elt-2::EGFP*. Scale bar, 50 μm.

The earlier appearance of *punc-119::GFP* expression is notable since embryonic expression of this reporter occurs earlier than the others, and was visible in embryos even before egg laying (Figure 3A). This suggests that the order of expression of embryonic markers might follow the same sequence as that in embryogenesis, consistent with recapitulation of embryonic programs within the tumor. To test this further we crossed *punc-119::mCherry* and *pelt-2::EGFP* reporters into the same strains, and confirmed the earlier expression of *punc-119::mCherry* (Figure 3C).

Notably, the different markers were expressed at different locations within tumors (Figure 3D). Red fluorescence from *punc-119::mCherry* was distinguishable from red autofluorescence, as the former was brighter and more widely distributed (Figure 3—figure supplement 1A). Furthermore, examination of fluorescent regions within embryos using bright field microscopy did not reveal the presence of embedded embryos, confirming that tissue derived from unfertilized oocytes was expressing embryonic genes (data not shown).

In mutationally-induced *C. elegans* germline teratomas, as in mammalian teratomas, germ cells transdifferentiate into various somatic tissues including intestine, muscle and neurons (Ciosk et al., 2006). It seemed to us unlikely that actual tissue differentiation occurs in wild-type uterine tumors, since these are formed from a small number of large, hypertrophic cells. To explore this we looked for the presence of several markers of tissue differentiation: blue autofluorescent gut granules (intestine), expression of *unc-119::GFP* in filamentous processes (neuron), and expression of *myo-3::GFP* in a striated pattern (muscle), but no case was differentiation detected. MYO-3::GFP fluorescence was sometimes seen in embedded embryos (Figure 3—figure supplement 1B). To verify that such fluorescence issued from embedded embryos rather than teratomas, we examined a fertilization defective *rrf-3(b26); edIs6 [unc-119::GFP + rol-6(su1006)]* strain. The presence of *rrf-3(b26)* (previously known as *fer-15(b26)*) largely abrogated MYO-3::GFP fluorescence within tumors (Figure 3—figure supplement 1C).

The status of germline chromatin prevents expression of somatic genes, and loss of the LIN-53 histone chaperone can promote neuronal differentiation within germline (Tursun et al., 2011). To probe the role of *lin-53* as a teratoma suppressor in uterine tumors, we tested effects of *lin-53* RNAi on tumor expression of an embryonic neuronal marker (*punc-119::GFP*). However, no effect of *lin-53* RNAi was detected (Figure 3—figure supplement 1D).

Overall, these results imply that uterine tumors to some extent recapitulate embryonic gene expression programs, but that this does not lead to full blown teratomas with embryonic differentiation, perhaps due to the absence of cell proliferation. Taken together with the etiological similarity to ovarian teratomas, this implies that uterine tumors are teratoma-like. An alternative possibility is that the disordered nature of chromatin masses leads to a broad deregulation of gene expression. However, the earlier expression of markers of early embryogenesis, and different location of expression of different embryonic markers argues against this interpretation.

### Tumor development is initiated by sperm depletion

To understand a disease, ideally one wants to identify its initial cause. For example, the original cause of malaria is the bite of an *Anopheles* mosquito that transmits protozoa of the genus *Plasmodium*. A long-standing question is: what is the origin of senescence? When and how exactly does it begin? Does such a starting point even exist? In the case of *C. elegans* uterine tumors, it is possible to provide a clear answer this question, as follows. Mature uterine tumors are a form of gross pathology that develops from normal oocytes, apparently as a consequence not of molecular damage but of quasi-programmed run-on of embryogenetic processes. Here, the precise moment of origin of pathology, equivalent to the mosquito's bite, may be defined: it is the moment after the last functional sperm in the spermatheca fertilizes an oocyte. This is the point of transition from program to quasi-program. Shortly thereafter, the first unfertilized oocyte enters the uterus and begins to undergo endomitosis, and the process begins that eventually produces a teratoma-like pathology.

This account suggests that it should be possible to accelerate or delay the development of uterine tumors by bringing forward or delaying sperm depletion, i.e. altering the timing of the mosquito bite (the transition from program to quasi-program). To test this we first examined *fog-2(q71)* mutants, which due to germline feminization do not produce sperm thereby changing hermaphrodites into females (Schedl and Kimble, 1988). Unfertilized oocytes accumulate in unmated *fog-2* worms, which have a very low ovulation rate producing only ∼20 oocytes before entering quiescence (Miller et al., 2003). Tumors were found to develop earlier in *fog-2* females than in N2 hermaphrodites (Figure 4A,B). The presence of tumors in *fog-2* females shows that sperm-derived signals such as the major sperm protein (MSP) are not required for tumor development. By contrast, tumor formation was suppressed by mating in *fog-2* females and N2 hermaphrodites (Figure 4A-C). Thus, altering the timing of the transition from program to quasi-program alters the rate of development of the senescent pathology that the quasi-program generates.

**Figure 4.**
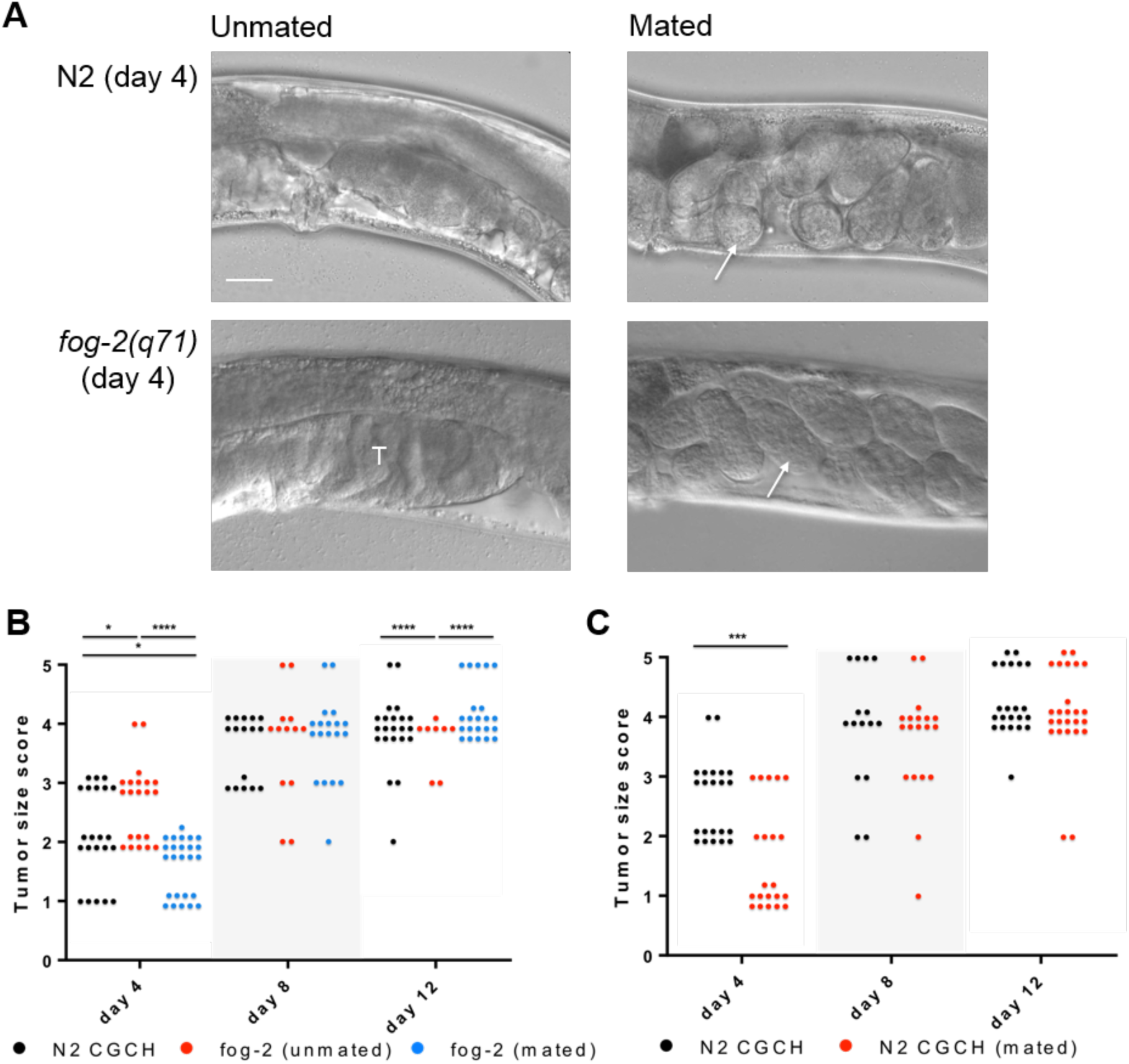
(A, B) Effects of *fog-2(q71)* and mating on timing of tumor development. (A) Selected images comparing N2 and *fog-2* tumors on day 4, and unmated and mated. Arrows, embryos; T, tumors. Scale bar, 25 μm. (B) Quantitation of tumor development. (C) Quantitation of tumor development in N2 hermaphrodites, unmated and mated. *, *p*<0.05; **, *p*<0.01; ***, *p*<0.001; ****, *p*<0.0001.

### Identification of genes contributing to tumor development

If uterine tumors result from embryogenetic quasi-programs, then this predicts that some genetic determinants of early embryogenesis will promote tumor development. To test this prediction we used RNA-mediated interference (RNAi) to knock down function of candidate genes, testing for suppression of tumor growth. Candidates included genes specifying germline development and the cell cycle (Figure 5—table supplement 1) (Gonczy et al., 2000; Korzelius et al., 2011; van den Heuvel, 2005). As positive controls, we includes two genes involved in protein synthesis (*rps-1*, *egl-45*), where RNAi was previously shown to inhibit germline development (Gonczy et al., 2000). Findings in the preceding section imply that RNAi-induced retardation of oocyte production rate could cause a delay in sperm depletion and consequently of the onset of tumor development. We therefore first tested RNAi initiated on day 3 of adulthood, to try to limit the effect of gene knockdown to the period after sperm depletion. The positive controls (*rps-1*, *egl-45*) both reduced tumor size, showing efficacy of RNAi within the uterus, but did not affect nuclear size (Figure 5A). Of other knockdowns, only that of *wee-1.3* reduced tumor size. However, in most cases a significant reduction of nuclear hypertrophy was seen.

**Figure 5.**
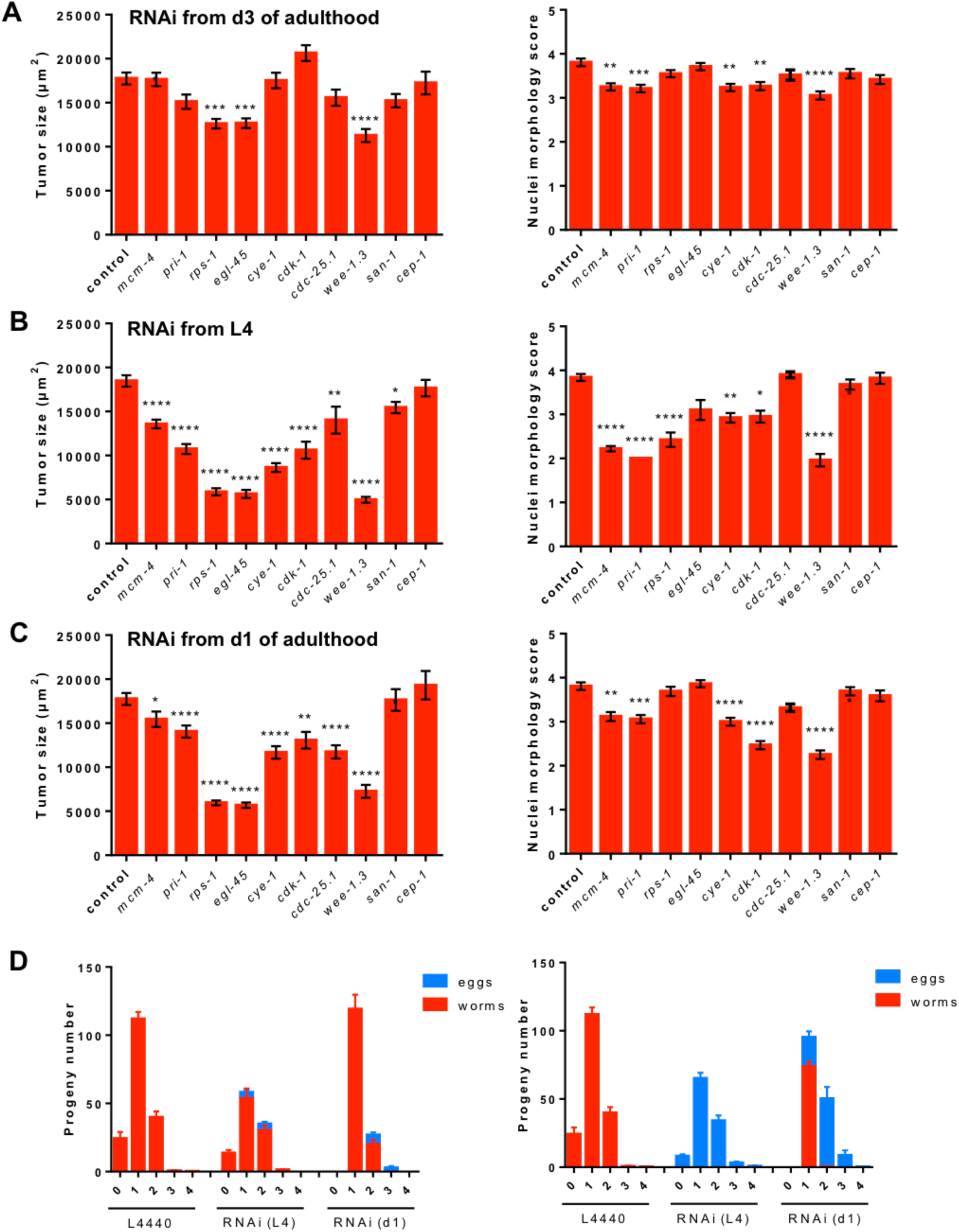
Knockdown of genes required for early embryogenesis reduces tumor growth. (A) Summary bar graph showing effects of RNAi from day 3 on nuclear hypertrophy and tumor size. (B) Summary bar graph showing effects of RNAi from L4 on nuclear hypertrophy and tumor size. (C) Summary bar graph showing effects of RNAi from day 1 on nuclear hypertrophy and tumor size. A., B., C., Tukey multiple comparison test, Wilcoxon-Mann Whitney test, *, *p*<0.05, **, *p*<0.01, ***, *p*<0.001, ****, *p*<0.0001. (D) RNAi effects on fertility schedule.

The lack of effects on tumor size could reflect weak reduction of gene function. We therefore tested RNAi initiated earlier, on L4 and day 1 of adulthood, and this resulted in major reductions in both tumor size and nuclear hypertrophy in most cases (Figure 5B, C). This included inhibition of genes involved in DNA replication (e.g., *mcm-4* and *pri-1*) and cell cycle promoters (e.g. *cye-1* and *cdk-1*). RNAi of *wee-1.3* also had similar effects, which was opposite to the expected result, since this gene encodes a CDK inhibitory kinase (van den Heuvel, 2005). Interestingly *wee-1.3* RNAi also resulted in a novel phenotype: development in the distal gonad of additional tumors formed, like uterine tumors, from a small number of large, hypertrophic cells with enlarged nuclei (Figure 5—figure supplement 1A). This suggests that loss of *wee-1.3* inhibition in the distal gonad leads to cellular hypertrophy. Given the negative correlation between the respective sizes of uterine tumors and the distal gonad (Figure 1—figure supplement 1E,F), we postulate that development of distal gonad hypertrophic tumors leads to reduced uterine tumor size, by the same mechanism. Again RNAi of the protein synthesis genes reduction of tumor growth but not of nuclear hypertrophy.

To check that reduction in tumor size were not merely the result of a delay in sperm depletion, we examined the effect of RNAi of productive profiles for selected genes (Figure 5D). RNAi of *cye-1* and *cdk-1* from L4 and day 1 did not delay the reproductive schedule, indicating that the timing of sperm depletion was unaffected. These results provide evidence that the function of wild-type genes specifying functions required for early embryonic development, including the cell cycle and protein synthesis, promote growth and reproduction in early life and senescent uterine tumor development after reproduction, another example of antagonistic pleiotropy as run on (Blagosklonny, 2006; Williams, 1957).

### Links between tumor growth and other senescent pathologies

This study presents evidence that run-on of embryogenetic mechanisms leads to uterine tumor formation. However, additional factors are likely to contribute to tumor growth, as follows. Prior to ovulation, developing oocytes import large quantities of yolk (vitellogenin and yolk lipid) from the body cavity using the LDL receptor-like protein RME-2 (Grant and Hirsh, 1999). Tumors in a *C. elegans* strain expressing a GFP-tagged vitellogenin, VIT-2::GFP, show strong green fluorescence (Figure 6A), suggesting that oocytes within the uterus continue to import yolk. Consistent with this, staining of neutral lipids with the fluorescent dye Bodipy showed high levels of lipid within tumors (Figure 6B).

**Figure 6.**
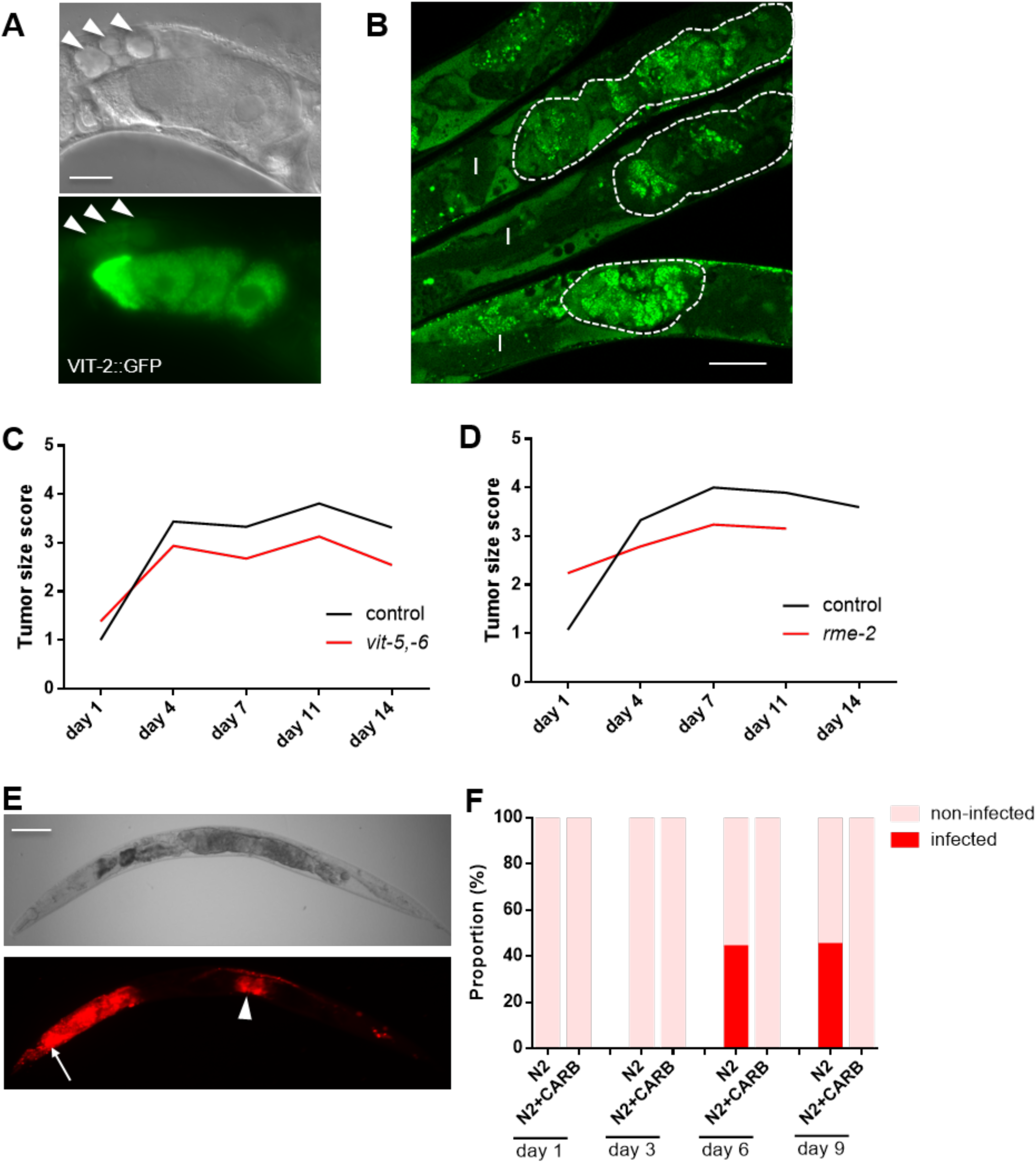
(A) VIT-2::GFP accumulation in uterine tumors (day 8 adult). Left, prior to sperm depletion. Note VIT-2::GFP in mature oocytes and embryos. Right, VIT-2::GFP accumulation in tumors. VIT::GFP in yolky pools (arrowheads). Scale bar, 25 μm. (B) Neutral lipid staining shows lipid accumulation within uterine tumors (day 8 adult, 15 μM FUDR). I, intestine. Scale bar, 25 μm. (C) *vit-5, −6* RNAi reduces tumor size. Summed data from 3 trials. (D) *rme-2* RNAi reduces tumor size. Summed data from 2 trials. (E, F) Uterine tumors can develop bacterial infections. (E) *E. coli* expressing RFP within uterine tumors. (day 10 adult, necropsy) Arrow, infected pharynx. Arrowhead, infected tumors. Scale bar, 100 μm. (F) Bar graph showing frequency of worms with infected tumors, without or with carbenicillin.

One possibility is that uptake of yolk contributes to tumor growth, either by increasing their bulk through its presence, or by providing nutrients to support new biosynthesis within tumor cells. To test this, we first used combined RNAi of *vit-5* and *vit-6*, which largely abrogates yolk accumulation in adults (Ezcurra et al., 2017), and found that it reduced tumor size (Figure 6C). RNAi of the *rme-2* yolk receptor had a similar effect (Figure 6D). Thus, yolk uptake by oocytes promotes tumor growth. This suggests that yolky lipid pools that accumulate in the body cavity (Ezcurra et al., 2017; Garigan et al., 2002) promote further pathology by feeding uterine tumors.

We also noted that wild-type *C. elegans* maintained upon *E. coli* expressing red fluorescent protein (RFP) sometimes develop strong red fluorescence within tumors (Figure 6E). This suggests that uterine tumors can become infected by the *E. coli* upon which *C. elegans* are routinely cultured. Examination of RFP positive tumors using light microscopy confirmed the presence of *E. coli* within the tumors (data not shown). One possibility is that such infections are nourished by the rich yolky milieu of the tumor. Such infections are prevented by treatment with the antibiotic carbenicillin (Figure 6F), revealing another mechanism by which antibiotics reduce senescent pathology and, possibly, extend lifespan (Garigan et al., 2002).

Under standard culture conditions some 40% of wild-type N2 hermaphrodites die as the result of pharyngeal infection with *E. coli* (Zhao et al., 2017). One possibility is that such worms have weaker organism-wide immunity which also leads to uterine tumor infection. To test this we examined worms on days 10 and 14, when P infections are relatively frequent, to see whether those with P had a higher frequency of infected tumors, but they did not, arguing against this (Figure 6—figure supplement 1). Thus, independent mechanisms lead to bacterial infection of the pharynx and uterus.

Taken together, these findings illustrate how distinct etiologies (embryogenetic and vitellogenic quasi-programs, and bacterial infection) act together in the genesis of a major senescent pathology.

## Discussion

A central claim of the Williams Blagosklonny theory is that quasi-programs promoted by late-life action of wild-type genes are a major cause of late-life pathology, including pathologies that limit lifespan. Previous studies have shown how several senescent pathologies of *C. elegans* result from quasi-programs. For example run-on of physiological apoptosis contributes to atrophy of the hermaphrodite gonad (de la Guardia et al., 2016), and run-on of yolk production leads to yolk steatosis and intestinal atrophy (Ezcurra et al., 2017; Herndon et al., 2002). Here we characterize in detail the developmental pathology of the large uterine tumors that are such a salient anatomical feature of aging *C. elegans*, and which provide a particularly clear example of how quasi-programs cause senescent pathology - i.e. of actual etiology in senescence. Importantly, this etiology is similar to that of human ovarian teratomas, demonstrating the conservation of quasi-programmed etiologies between *C. elegans* and humans. This, despite the outward differences between worm and human pathology.

### Cell cycle run-on causes uterine tumor development

This study supports the view that uterine tumors result from an embryogenetic quasi-program, which initially manifests as run-on of endomitosis. Active DNA replication in DNA masses has been previously demonstrated using 5-bromo-2-deoxyuridine (BrdU) incorporation (Golden et al., 2007). This leads to polyploidy, nuclear hypertrophy and cellular hypertrophy. Increased ploidy promotes growth in many contexts, e.g. in the *C. elegans* hypodermis (Lozano et al., 2006), and we postulate that by similar mechanisms increased oocyte ploidy causes cellular hypertrophy.

In many cell types DNA synthesis in the absence of karyokinesis and cytokinesis is prevented by the G1 and G2 cell cycle checkpoints. However, in a number of species these checkpoints are absent in early embryogenesis. This absence, which promotes embryogenesis also promotes tumor formation.

Endomitosis in unfertilized oocytes may be supported by maternal transcripts intended to support zygotic cell cycling (Iwasaki et al., 1996). Our observations of expression in older tumors of genes expressed during embryogenesis suggests action of maternal transcripts in cellular differentiation too.

Runaway endomitosis in unfertilized *C. elegans* oocytes does not occur in unfertilized eggs in *Drosophila* thanks to the presence of an arrest point after meiosis II in the latter, defects in which can result in giant polyploid nuclei within eggs (Doane, 1960), as seen in *C. elegans*. This potentially provides insight into the evolution of longer life: since female *Drosophila* may take some time to be mated, they need to maintain a stock of viable unfertilized eggs, which leads to selection for the post-meiosis II arrest point. By contrast *C. elegans* mature oocytes are immediately fertilized by self sperm, which is predicted to weaken selection for such an arrest point.

In mammals, cancer is considered to be primarily the consequence of somatic mutations affecting the control of cell proliferation. However, the age-increase in cancer rate is to some degree the result of the effect on aging on the cellular microenvironment (Campisi, 2013). Moreover, some forms of cancer can originate with hyperfunction rather than mutation, for example benign prostatic hyperplasia in humans, which typically precedes prostate cancer, is the result of long-term testosterone exposure rather than mutation (Waters et al., 2000). Here we postulate that the primary cause of uterine tumors is an embryogenetic quasi-program. An additional possibility is that the resulting chromatin hypertrophy triggers a DNA damage response, with further pathogenetic effects (McGee et al., 2012).

### Tumor growth will affect expression profiles in aging *C. elegans*

Given the large size of uterine tumors, their development likely contributes to age-changes in whole worm expression profiles (where the germline is present). For example, one study using fertilization-defective mutants (e.g. *rrf-3(b26)*, at 25°C) found that many oocyte-expressed mRNAs increase in abundance after day 3 of adulthood, and remain highly abundant until late life (Lund et al., 2002). Similarly, protein profiles from aging *C. elegans* (Walther et al., 2015) exhibited an age increase in germline associated proteins that remained high long after the end of reproduction (Walther et al., 2015). We postulate that this reflects the increase in histone in the DNA masses of uterine tumors, consistent with the large increases in HIS-58::GFP expression that we have observed (Figure 2A,C,D and Figure 2—figure supplement 2C). Proteins whose levels increase in aging hermaphrodites (FUDR-treated) also include five abundant ULE (uterine lumen expressed) proteins of unknown function (Zimmerman et al., 2015).

Such age-changes in mRNA and protein are more likely the consequence than the cause of pathogenic processes such as run-on of embryogenetic function in unfertilized oocytes, particularly run-away endomitosis.

### Wild-type genes promoting embryogenesis generate pathology

Knockdown of genes required for embryogenesis, including cell cycle genes, inhibits tumor growth. These findings are consistent with previous observations that blocking DNA replication using FUDR is sufficient to prevent formation of chromatin masses formation (Golden et al., 2007) and uterine tumors (Riesen et al., 2014). They demonstrate that wild-type genes encoding cell cycle machinery are pathogenic in later life, and illustrate enactment of antagonistic pleiotropy by quasi-programs (Blagosklonny, 2006; Williams, 1957). Here the initial cause of senescent pathology is not damage or loss of homeostasis, but the late-life action of wild-type genes and the biological processes that they specify.

### Similarities between uterine tumors and mammalian teratomas

*C. elegans* uterine tumors are pathophysiologically similar to mammalian ovarian teratomas. For example, both are tumors derived not from mutation, but from run-on of embryogenetic programs in unfertilized oocytes. In the case of *C. elegans* this gives rise to amorphous masses derived from hypertrophic polyploid oocyte-derived cells. They show expression of genes associated with later embryonic development, but no detectable morphological differentiation, and no cell proliferation, in contrast to mammalian teratomas. Thus, *C. elegans* uterine tumors are teratoma-like. A further similarity is that human ovarian teratomas are usually benign, and *C. elegans* uterine tumors do not increase late-life mortality (Riesen et al., 2014).

A difference between *C. elegans* uterine tumors and mammalian ovarian teratomas is that the former is a senescent pathology but not the latter, which instead occur largely during reproductive years (in humans between ∼20-40 years of age) (Ayhan et al., 2000). However, the fact of the etiological features that they share - in broad terms, quasi-programs resulting from wild-type gene function - yields a dark insight into the nature of aging: that senescent pathology and teratoma are to some extent pathophysiologically equivalent. That is to say: insofar as senescent pathology is caused by quasi-programs, aging and teratoma are the same sort of disease. By the same token, cholera and tuberculosis are similar sorts of disease, insofar as both are caused by bacterial infection.

### Is the uterus a latent tumor niche?

It is notable that in post-reproductive hermaphrodites, unused oocytes remain in both the uterus and the proximal gonad, but only the former develop into tumors. Why might this be? One clue is that the presence of oocytes within the uterus is not programmed but quasi-programmed. It was previously shown that if mitotic germ cells are mislocated near the somatic gonadal sheath, the latter acts as tumor niche, causing hyperplastic proximal tumors (the Pro phenotype). It has been demonstrated that Notch ligand expression creates a latent tumor niche at the somatic gonad (McGovern et al., 2009). Moreover, the uterus secretes a number of proteins of unknown function into its lumen (Zimmerman et al., 2015). This raises the question: could the uterus also be a latent tumor niche? In a preliminary screen of notch ligand gene knock-down and mutation, we observed that *dsl-5(ok588)* slows tumor growth without affecting final tumor size or altering fertility (data not shown). However, the site of expression of *dsl-5* remains unclear (McGovern et al., 2009).

## Multiple etiologies of tumorigenesis

Uterine tumors illustrate how different etiologies combine to promote senescent pathology. Here the initial etiology is run-on of embryogenetic quasi-programs. Subsequently, a second pathology, yolk steatosis, contributes to tumor growth, in a distant parallel to the increased cancer risk caused by obesity in mammals. Thirdly, tumors can become infected with bacteria, and here again there are resemblances to mammalian pathogenesis, as follows.

The relationship between bacteria and cancer progression in humans is a long-standing topic of investigation. In some instances, bacteria appear to initiate or drive malignant alterations. For example, colorectal cancer is often associated with an abundance of *Streptococcus gallolyticus* (formerly *S. bovis*) and *Fusobacterium sp*, which are thought to trigger and drive cancer progression by causing persistent inflammation (Cummins and Tangney, 2013; Kostic et al., 2012). Here the anaerobic conditions and available nutrients within tumors provide an optimum environment for the growth of certain bacteria (Cummins and Tangney, 2013). Notably, bacterial infection of tumors is not necessarily detrimental to the host. Post-operative infection in lung cancer can actually improve patient survival (Ruckdeschel et al., 1972), perhaps because bacteria compete for nutrients, secrete toxic proteins or help to initiate immune responses against tumor cells. Given that certain bacterial species have an affinity for particular cancers (Cummins and Tangney, 2013), genetically engineered bacteria are now being developed as potential anti-cancer therapies (Baban et al., 2010). These bacteria can either secrete tumor-suppressor proteins or act as vectors to deliver therapeutic genes via plasmids to cancerous cells (Baban et al., 2010). Given the susceptibility of *C. elegans* uterine tumors to bacterial invasion as demonstrated here, they might serve as a model to study this tumor-bacteria interactions, or to test bacteria-based anti-cancer therapies.

## Conclusions

The main purpose of studying aging is *C. elegans* is to understand senescence, particularly in terms of its original causes. This is part of the broader, general goal of biomedical research: to understand the etiologies of disease in order to be able to prevent and treat them (Gems, 2015). This study describes the primary mechanistic cause of one component of *C. elegans* aging, the uterine tumors. It exemplifies a class of diseases of aging caused by quasi-programs that form a major part of the *C. elegans* aging process. Thus, this work brings us closer to a mature understanding of the nature of aging in *C. elegans*. We postulate that in *C. elegans*, quasi-programs are the major cause of senescence, along with other contributory causes, such as mechanical senescence and bacterial infection (Podshivalova et al., 2017; Zhao et al., 2017)(this study). It seems likely that in human aging molecular damage, particularly DNA damage, plays a greater role than in *C. elegans*.

## Materials and methods

### *C. elegans* culture and strains

Worms were cultured at 20°C unless otherwise stated using standard method described previously (Brenner, 1974). Nematode strains used in this paper include N2 (wild-type) CGCH (Gems and Riddle, 2000), CB4108 *fog-2(q71) V*, DP132 *edIs6 [unc-119::GFP + rol-6(su1006)] IV*, GA1932 *unc-119(ed3) III; ltIs44 [pie-1p-mCherry::PH(PLC1deltal)+unc-119(+)]; ruIs32 [pie-1::GFP::H2B+unc-119(+)] III*, OP37 *unc-119(ed3) III; wgIs37 [pha-4::TY1::EGFP::3xFLAG + unc-119(+)]*, OP56 *unc-119(ed3) III; gaIs290 [elt-2::TY1::EGFP::3xFLAG(92C12) + unc-119(+)]*, OP64 *unc-119(ed3) III; wgIs64 [hlh-1::TY1::EGFP::3xFLAG + unc-119(+)]*, OP354 *unc-119(tm4063) III; wgIs354 [elt-1::TY1::EGFP::3xFLAG + unc-119(+)]*, PS3662 *syIs63[cog-1::GFP + unc-119(+)]*, and RW10006 *ruIs32* [*ppie-1::histoneH2B::GFP*] *zuIs178* [*phis-72::his-72::GFP unc-119(+)*] *unc-119(ed3) III.*

### Mating protocol

N2 or *fog-2(q71)* males were used for mating. Sexes were separated at L4 and allowed to develop into adults before being combined for mating. A brief mating protocol was used to reduce the life shortening effects of mating, as described (Gems and Riddle, 1996).

### Microscopy

Nematodes were places on 2% agarose pads and anesthetized with 0.2% levamisole. Nomarski and epifluorescence microscopy was performed on a Zeiss Axioskop 2 Plus microscope connected to a Hamamatsu C10600 - Orca ER digital camera. Images were acquired and quantified using Volocity 6.3 software. 100x or 400x images are shown in this report. For viewing red autofluorescence, a rhodamine filter was used, l_ex_ 545/25 nm, l_em_ 605/70 nm).

Selective plane illumination microscopy (SPIM) was used to make 3D reconstructions of *C. elegans* uterine tumors. Anaesthetized worms were transferred into a plastic capillary tube with 1.5% low melting point agarose, 0.03% levamisole and 0.5 mm sized FluoSphere microspheres (beads) (1:1000, F8813 from Life Technologies). Agarose embedding and anesthetic kept the animals immobile during imaging, and FluoSphere beads allowed registration and 3D construction of 2D images taken from 5 angles. We used a 1 ml BD Plastikpak (REF 300013) syringe to mount the plastic capillaries into the OpenSPIM chamber full of M9 buffer. OpenSPIM was performed to take images as previously described (Girstmair et al., 2016). Acquired data was processed using Fiji software. The beads registration algorithm and the multi-view deconvolution plugin were performed for reconstruction of 3D structure (Preiblsh et al., 2010; Preibls et al., 2014).

### DAPI staining and quantitation of genomic copies

Nuclei of uterine tumors were stained with the DNA-binding dye 4’, 6’-diamidino-2-phenylindole (DAPI). Worms of different ages were fixed with methanol and incubated on ice, then washed with M9 buffer and stained with 500 ng/ml DAPI staining in darkness for 30 min. Finally, worms were washed again with M9 buffer before imaging. Genomic copies were quantified using real-time PCR as described (Golden et al., 2007).

### Pathology scoring system

Severity of uterine tumors were scored using a five stage classification as previously described (Riesen et al., 2014). Score 1 denotes a uterus containing fertilized eggs or oocytes of normal size and morphology. Score 2 denotes a uterus containing unfertilized oocytes only, with a slightly abnormal appearance. Score 3 and score 4 denote a small tumor and a large tumor, respectively. Score 5 denotes a very large tumor which fills the body cavity in mid-body region.

To measure different levels of nuclear hypertrophy we created another five stage classification. Score 1 denotes small, spherical early stage oocyte nuclei. Score 2 denotes larger but still spherical nuclei. Score 3 denotes larger nuclei with irregular morphology. Score 4 denotes highly hypertrophic nuclei with grossly abnormal phenotype (e.g. with major protrusions), such that that some individual nuclei are barely distinguished. Score 5 denotes large chromatin masses where most individual nuclei can no longer be distinguished.

To describe expression of embryonic reporters at different levels we used a four stage classification system. Score 1 denotes no reporter fluorescence in the tumor. Score 2 denotes weak reporter fluorescence in the tumor. Score 3 denotes strong reporter fluorescence in the tumor.

### RNA-mediated interference

The RNAi clones used in this paper were obtained from Ahringer library. Sequencing was used to confirm the identity of inserts of all these RNAi clones. RNAi treatment was performed as described previously (Kamath et al., 2001). Worms were fed with *E. coli* HT115 and transferred to plates seeded with RNAi clone at L4 stage, day 1 and day 3 seperately. *E. coli* HT115 containing empty vector L4440 was used as control.

### Statistical analysis

Correlation analysis was performed using Linear Regression analysis. Non-parametric Wilcoxon-Mann Whiney test was performed to compare tumor size and nuclear morphology. Fluorescence intensity was measured using Volocity and Fiji software. Multiple comparison t test was used to compare fluorescence intensity and tumor size.

## Supplemental information

Supplemental Information includes fives figures, two movies and two tables and can be found with this article online.

## Author contributions

D.G., H.W. and Y.Z. conceived the study and designed experiments. T.A. performed the spermathecal marker tests. A.F.G. characterized tumor infections. J.G. and M.T. assisted with SPIM microscopy. J.H. performed analysis of tumor development in animals with altered sperm availability. D.I. and J.T. assisted with analysis of profiling data. Other experiments were performed by H.W., Y.Z. and M.E.. D.G., H.W. and Y.Z. wrote the manuscript

## Acknowledgments

We thank Ryan Baugh and Jim Jordan (Duke University), Fadri Martinez-Perez (Imperial College), Richard Poole (UCL), and members of the Gems lab for useful discussion and/or comments on the manuscript. We also thank Stuart Kim (Stanford University) for making available microarray data. Some strains were provided by the Caenorhabditis Genetics Center, which is funded by NIH Office of Research Infrastructure Programs (P40 OD010440). This work was supported by a Wellcome Trust Strategic Award (098565/Z/12/Z) and an EU grant (FP6-518230) to D.G. and European Research Council (ERC-2012-AdG 322790) and Biotechnology and Biological Sciences Research Council (BBS/H006966/1) grants to M.T.

---

**Figure 1—figure supplement 1.**
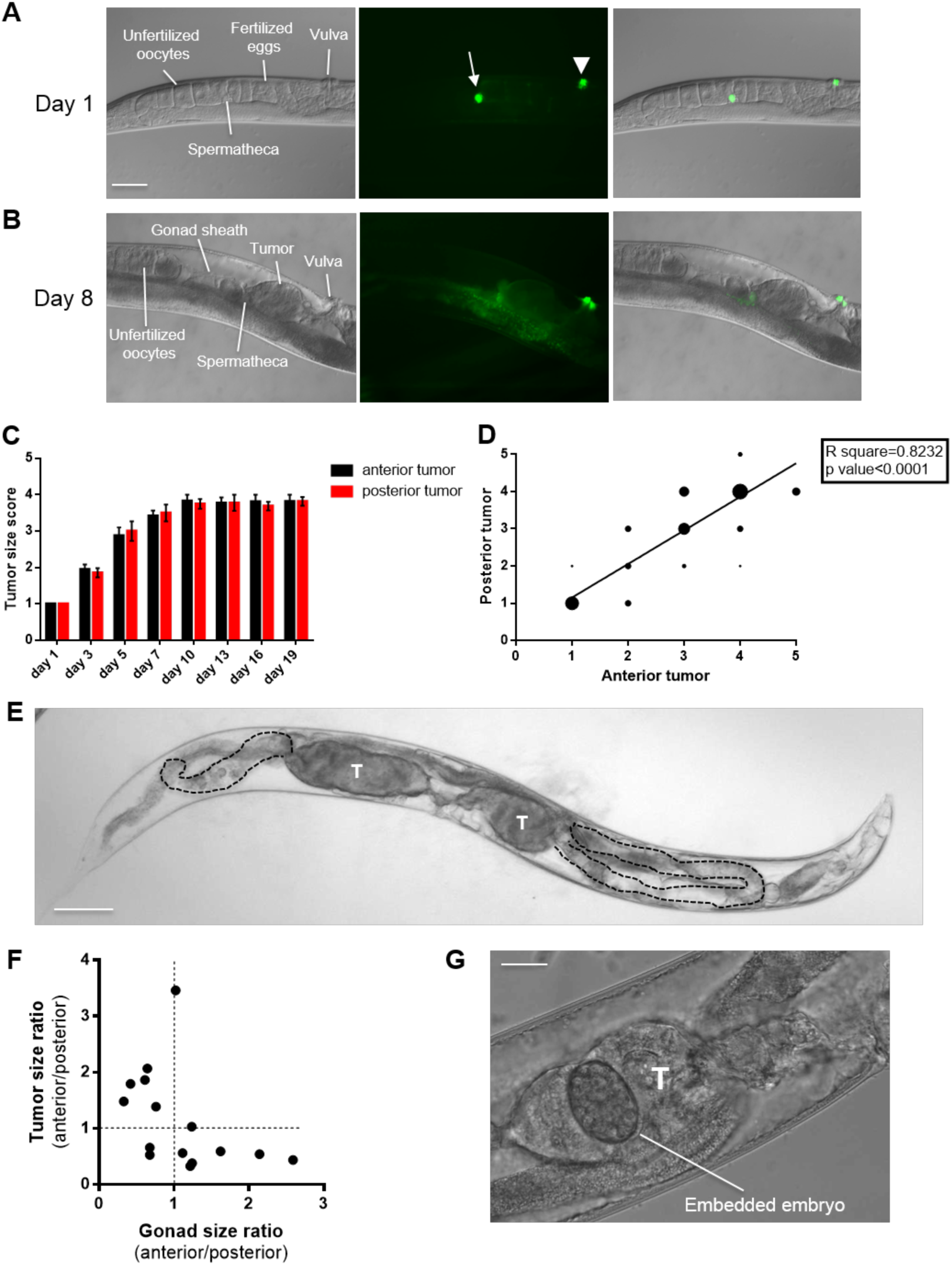
(A,B) Tumors are contained within the uterus. (A) Day 1 of adulthood. Normal gonad with no tumor, and GFP expression at spermathecal uterine valve and vulva. (B) Day 8, GFP expression at spermathecal uterine valve indicates that the distal end of the tumor is within the uterus. Arrow, spermatheca; arrowhead, vulva. (C) Anterior uterine tumors are not larger or smaller overall than posterior uterine tumors. (D) Correlation between size of anterior and posterior tumors. Linear regression test, *p*<0.0001. (E,F) Asymmetry in gonad disintegration in animals with asymmetrical uterine tumor pairs. (E) Day 19 adult with asymmetric tumors. Scale bar, 100 μm. T, tumor. (F) Ratio of anterior tumor/posterior tumor size and gonad size within the same worm with asymmetric tumors. (G) Embedded embryo within uterine tumor. (day 8 adult) Scale bar, 25 μm.

**Figure 1—figure supplement 2.**
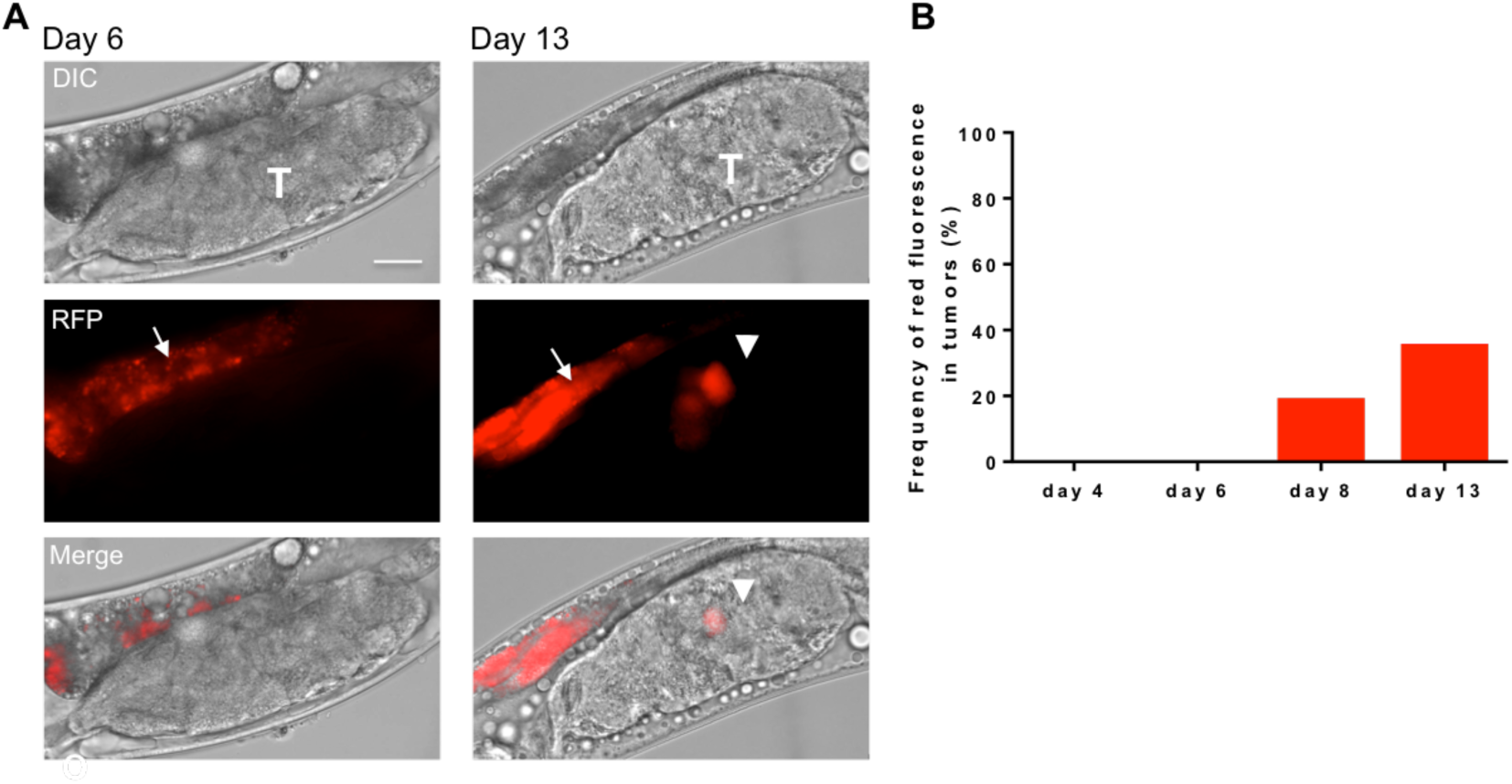
(A, B) Red fluorescence in older uterine tumors. (A) Red autofluorescence in tumor on day 13 (right) but not day 6 (left). T, tumor. Arrow, intestinal autofluorescence. Arrowhead, tumor autofluorescence. Strain, N2 CGCH. Scale bar, 25 μm. (B) Frequency of red fluorescence in uterine tumors with age. day 4, n=48, day 6, n=60, day 8, n=59, day 13, n=57.

**Figure 2—figure supplement 1.**
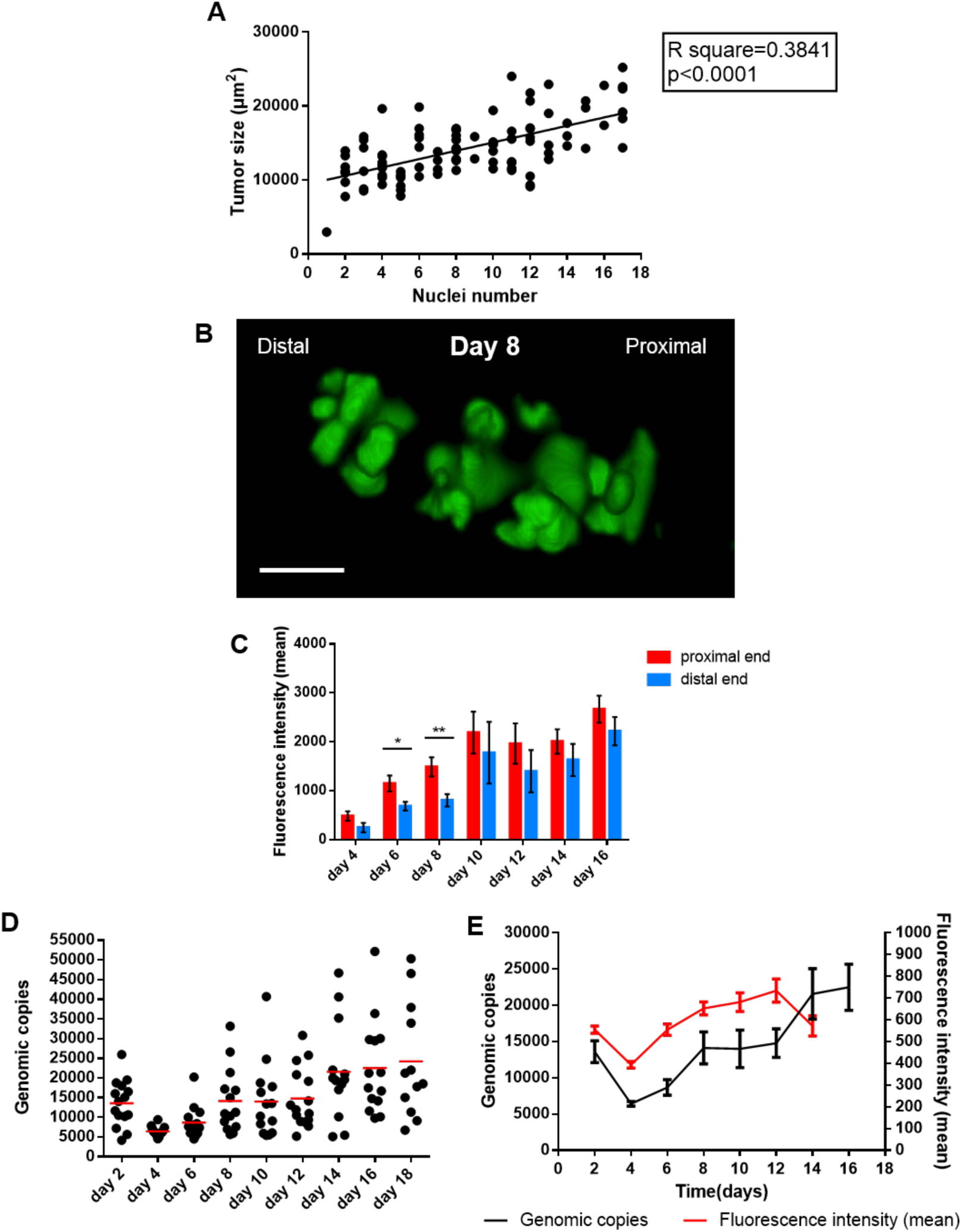
(A) Linear regression analysis of number of oocytes and tumor size. ****, *p*<0.0001. (B,C) Gradient hypertrophy from proximal end to distal end. (B) Tumour of day 8 adult (SPIM microscopy). Scale bar, 15 μm. (C) GFP fluorescence intensity of tumors. (D) Genomic copies at different days. Strain, GA1932 *unc-119(ed3) III; ltIs44 [pie-1p-mCherry::PH(PLC1deltal)+unc-119(+)]; ruIs32 [pie-1::GFP::H2B+unc-119(+)] III*. (E) Changes of overall genomic copies and GFP fluorescence with age.

**Figure 2—table supplement 1.**
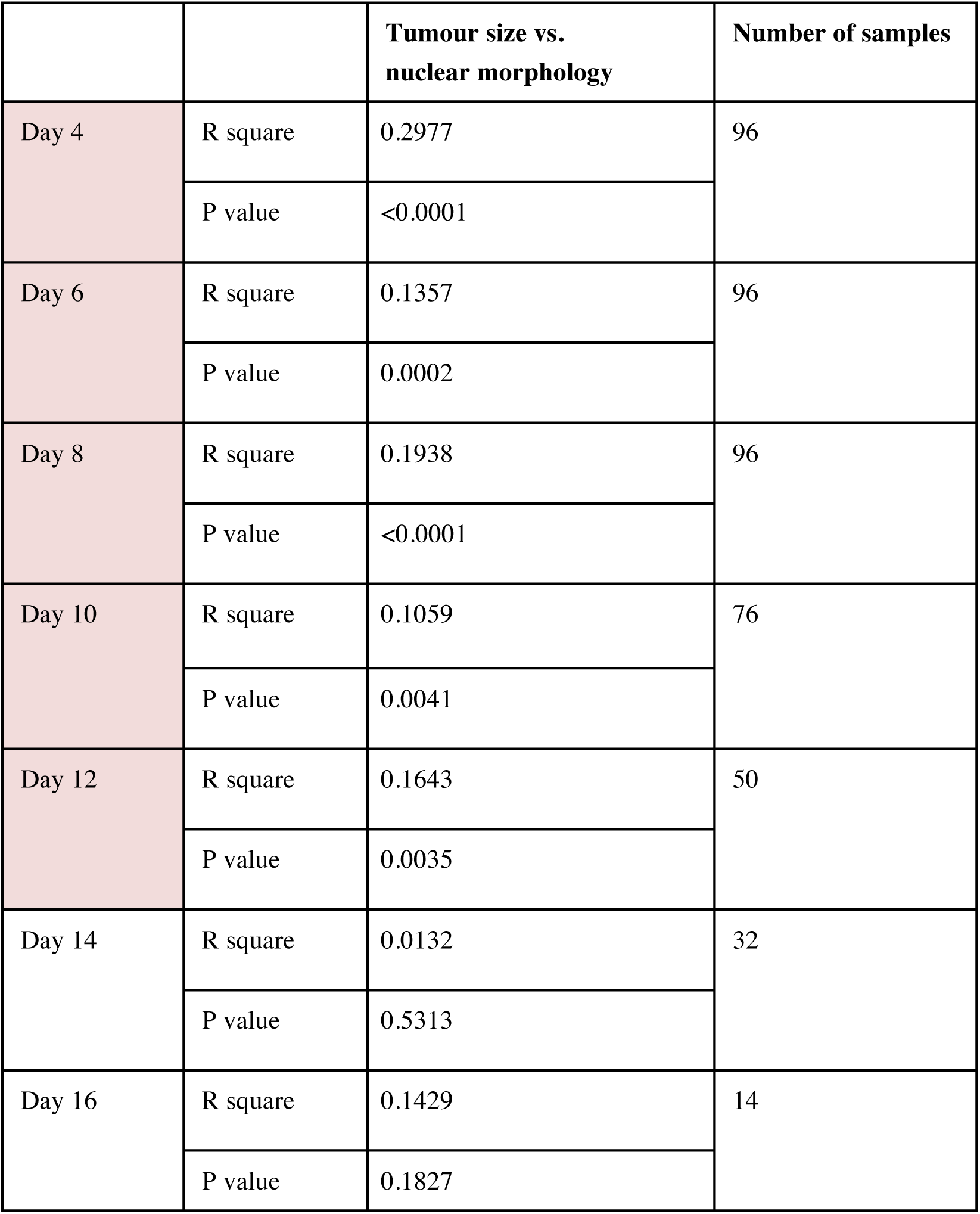
Correlation of tumor size and nuclear morphology on different days.

**Figure 3—figure supplement 1.**
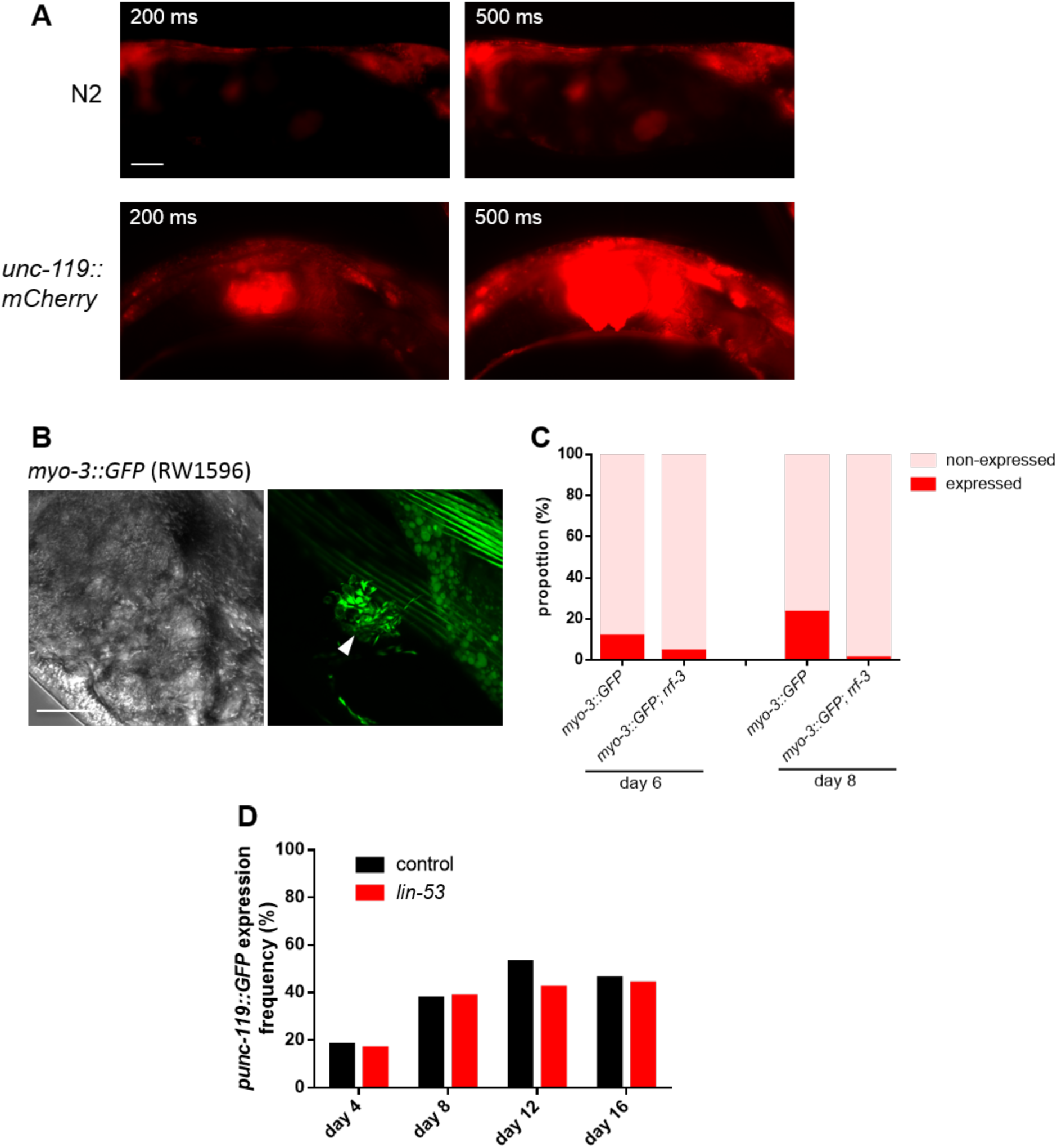
(A) Selected images of presence of red autofluorescence in N2, and mCherry fluorescence in *unc-119::mCherry* (MS1180) using different exposure time. Scale bar, 25 μm. (B) Tumor of day 8 adult with *myo-3::GFP* inclusion (Confocal microscopy). Scale bar, 25 μm. (C) Comparison of frequency of MYO-3::GFP positive inclusions in uterine tumor in *rrf-3(+)* and rrf-3(b26) strains. Rare, residual MYO-3::GFP positive inclusions in the latter resemble embedded embryos, suggesting possible incomplete penetrance of *rrf-3(b26)*. However, no larvae or dead eggs were seen on the NGM plates. (D) Effect of *lin-53* on frequency of tumors expressing *punc-119::GFP*.

**Figure 5—figure supplement 1.**
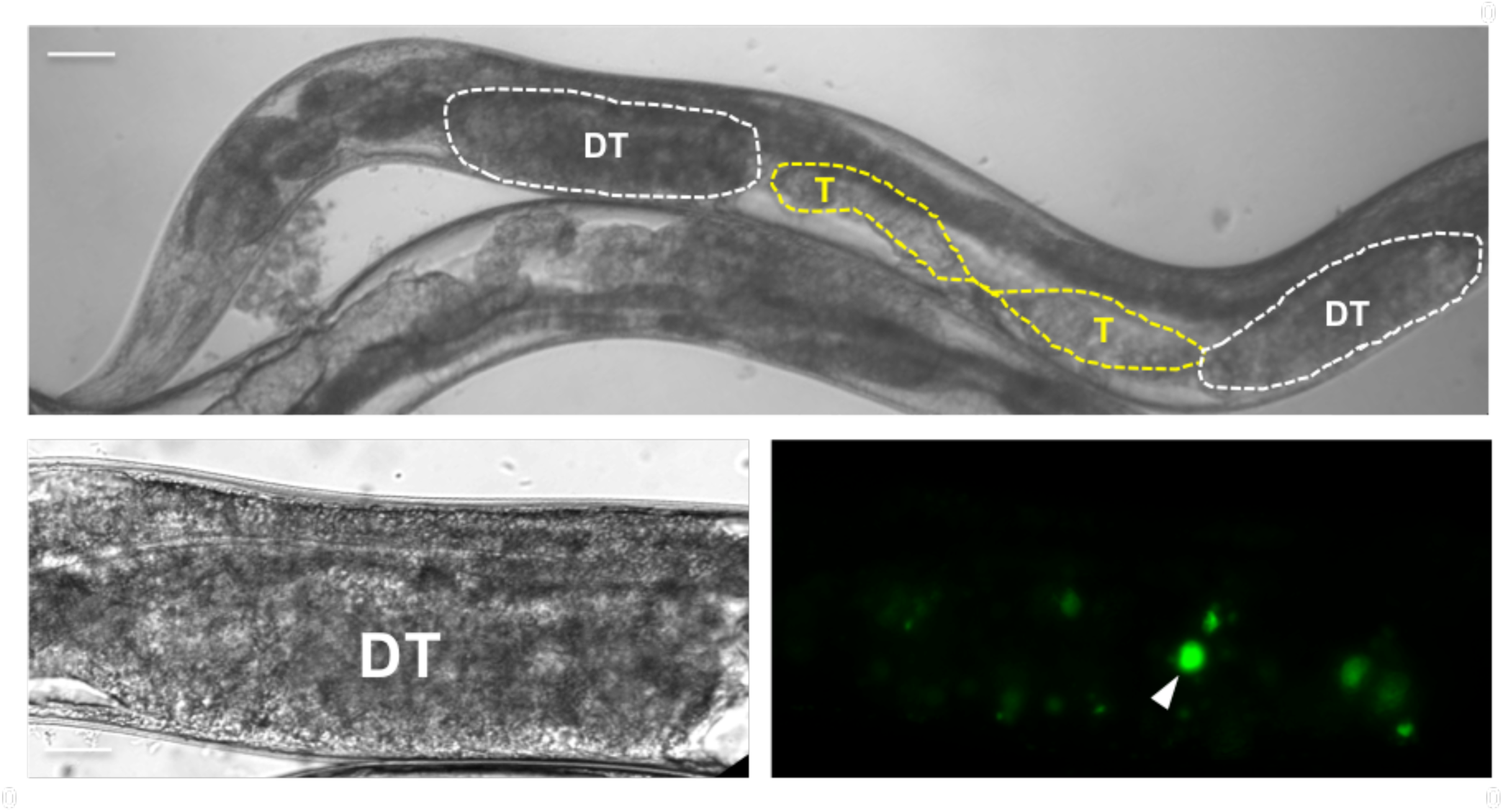
Distal gonad tumors resulting from *wee-1.3* RNAi (CDK inhibitory kinase). Bottom right: The size and number of HIS-58::GFP-labelled nuclei indicate that these tumors, like uterine tumors, result from cellular hypertrophy rather than hyperplasia. Arrowhead, hypertrophic nucleus. T, uterine tumor; DT, distal gonad hypertrophic tumor.

**Figure 5—table supplement 1.**
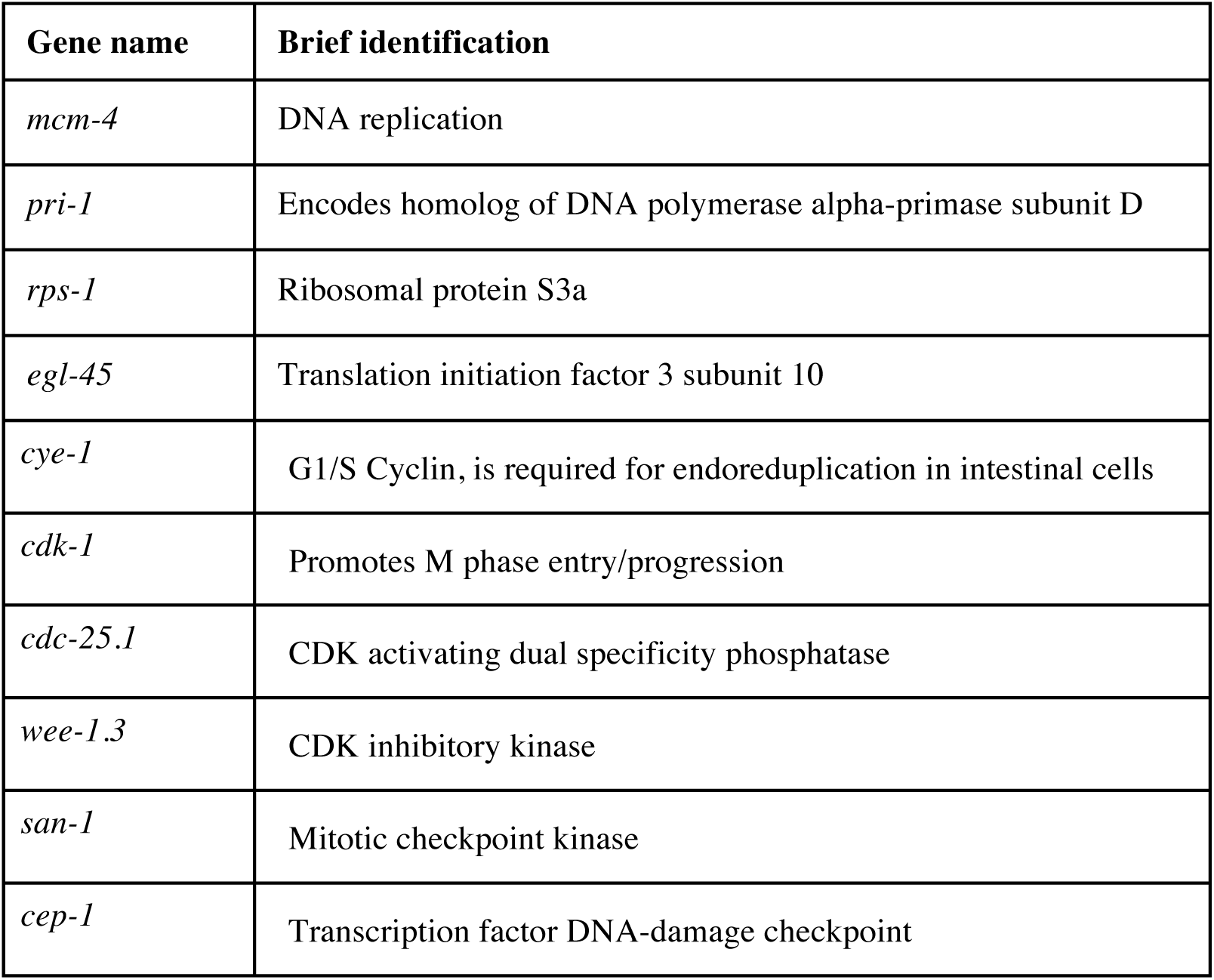
Gene list for RNAi experiment.

**Figure 6—figure supplement 1.**
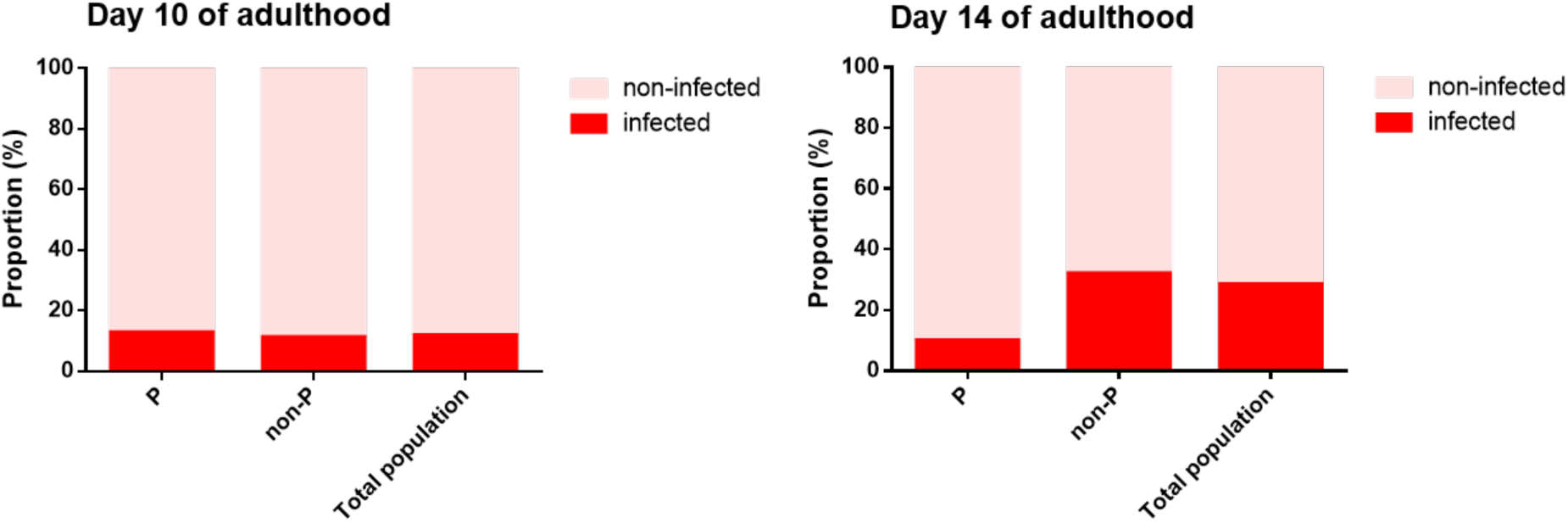
Worms with pharyngeal infection do not show a higher frequency of infected uteri. P, pharynx infected with RFP-tagged *E. coli*; non-P, not infected. The N2 line here used was CGCH (Gems and Riddle, 2000).

